# Investigating the ability of astrocytes to drive neural network synchrony

**DOI:** 10.1101/2022.09.26.508928

**Authors:** Gregory Handy, Alla Borisyuk

**Affiliations:** Departments of Neurobiology and Statistics, University of Chicago, Chicago, Illinois, USA; Grossman Center for Quantitative Biology and Human Behavior, University of Chicago, Chicago, Illinois, USA; Department of Mathematics, University of Utah, Salt Lake City, Utah, USA

## Abstract

Recent experimental works have implicated astrocytes as a significant cell type underlying several neuronal processes in the mammalian brain, from encoding sensory information to neurological disorders. Despite this progress, it is still unclear how astrocytes are communicating with and driving their neuronal neighbors. While previous computational modeling works have helped propose mechanisms responsible for driving these interactions, they have primarily focused on interactions at the synaptic level, with microscale models of calcium dynamics and neurotransmitter diffusion. Since it is computationally infeasible to include the intricate microscale details in a network-scale model, little computational work has been done to understand how astrocytes may be influencing spiking patterns and synchronization of large networks. We overcome this issue by first developing an “effective” astrocyte that can be easily implemented to already established network frameworks. We do this by showing that the astrocyte proximity to a synapse makes synaptic transmission faster, weaker, and less reliable. Thus, our “effective” astrocytes can be incorporated by considering heterogeneous synaptic time constants, which are parametrized only by the degree of astrocytic proximity at that synapse. We then apply our framework to large networks of exponential integrate-and-fire neurons with various spatial structures. Depending on key parameters, such as the number of synapses ensheathed and the strength of this ensheathment, we show that astrocytes can push the network to a synchronous state and exhibit spatially correlated patterns.

**Author summary:** In many areas of the brain, glial cells called astrocytes wrap their processes around synapses – the points of contact between neurons. The number of wrapped synapses and the tightness of wrapping varies between brain areas and changes during some diseases, such as epilepsy. We investigate the effect that this synaptic ensheathment has on communication between neurons and the resulting collective dynamics of the neuronal network. We present a general, computationally-efficient way to include astrocytes in neuronal networks using an “effective astrocyte” representation derived from detailed microscopic scale models. The resulting hybrid networks allow us to emulate and observe the effect of ensheathment conditions corresponding to different brain areas and disease states. In particular, we find that it makes the networks more likely to switch into a highly correlated regime, contrary to predictions from the traditional neurons-only view. These results open a new perspective on neural network dynamics, where our understanding of conditions for generating correlated brain activity (e.g., rhythms associated with various brain functions, epileptic seizures) needs to be reevaluated.

## Introduction

Recent theoretical work on neural circuits has successfully begun linking network structure to their underlying dynamics [1]. In particular, it has been shown that the statistics of connectivity (e.g., spatial scales of recurrent connectivity) can be used to predict the statistics of spiking behavior (e.g, level of network correlation) [2–4]. As reconstruction methods such as electron microscopy start to provide considerable amounts of data pertaining to the connectome of extensive brain regions [5–7], such theoretical work can guide the analysis of these massive datasets, suggesting the most useful network statistics to examine first. In this work, we develop a framework that extends computational spiking networks to efficiently include astrocytes, a non-neuronal cell that is often included in such reconstructed volumes [8]. As a result, this method allows for improving the scope and accuracy of theoretical predictions linking network structure to network dynamics.

While the overwhelming majority of computational networks only account for conventional synaptic connections across excitatory and inhibitory populations of neurons, many synapses found *in vivo* are ensheathed, or tightly wrapped around, by astrocytes [9–11]. It is well-known that astrocytes are engaged in bidirectional communication with nearby neurons, with recent breakthrough results demonstrating that astrocytes reliably encode sensory information [12]. Such communication is thought to occur through multiple pathways, such as neurotransmitter removal from the synaptic cleft, altering of extracellular ion concentrations, and the release of neuroactive substances [13–15]. The tightness of the ensheathment modulates the amount of extracellular space available around the synaptic site, and, as a result, is likely one of the contributing factors determining the properties of the synaptic connections [16] and the overall dynamics of the network. However, it is unclear how one may explore these factors computationally, as the field currently lacks an efficient method to implement such micro-scale details into a large spiking network.

Further, the presence of ensheathed and non-ensheathed synapses in a network would require a higher degree of heterogeneity than considered previously, since modeling efforts have largely focused on networks with weak heterogeneities across key synaptic parameter values to none at all [17–20]. Here, synaptic parameters would need to be drawn from distinct distributions depending on the presence of an astrocyte. Interestingly, it has been experimentally shown that the number of synapses ensheathed by astrocytes differs dramatically between brain regions (e.g., 74% of cerebellar Purkinje cell synapses, 29% of the dendritic spines in the mouse visual cortex, and 60% of hippocampal synapses are ensheathed [9, 10, 21]). Further, disease states such as epilepsy and Alzheimer’s disease are associated with astrocytes altered in a variety of ways (so called “reactive astrocytes”) [22–27], including alteration of their proximity to the synapse. Taken together, synaptic ensheathment by astrocytes is a source of heterogeneity in synaptic connections that needs to be accounted for to make possible predictions as to how neuronal dynamics shift across brain regions and states. Interestingly, this concept has not been included in previous mathematical studies that have investigate the so-called “tripartite synapse” (pre- and post-synaptic terminals plus neighboring astrocyte, choosing to assume the astrocyte portion does not vary from synapse to synapse [28–31].

Despite the wealth of evidence for synaptic ensheathment, the role it might play in regulating brain activity is unknown. For example, enhanced synaptic ensheathment during epilepsy in hippocampal CA3-CA1 network might be contributing to the run-away excitability and to seizures, or, conversely, they might be playing a compensatory, protective role [32]. We use computational modeling at different spatial scales to differentiate between these two options, and, more generally, to take a step towards understanding the role of astrocyte ensheathment in altering network dynamics.

As mentioned above, astrocytes have many pathways of interacting with neurons. In this work we limit ourselves to only considering two such interaction pathways, namely, the physical barrier around the synapses presented by astrocyte wrapping and a rapid removal of neurotransmitter from the synaptic cleft by glutamate transporters on the surface of astrocyte. Building on our earlier work on detailed synaptic models [33, 34], we show that the ensheathed synapse can be thought of as faster and weaker than the same synapse without tight wrapping by an astrocyte. This allows us to create an “effective astrocyte ensheathment” model, which can be included in established neuronal network frameworks with only minimal increases in computational cost. Namely, an “effectively ensheathed” synapse is endowed with parameter values (synaptic time constant and synaptic strength) derived from detailed studies of a synapse at the microscopic level. These synaptic properties are considered as parameterized only by the degree of astrocytic proximity at that synapse. This approach allows us to reevaluate several important examples of neuronal networks in which the dynamics had been well-studied and understood in the absence of astrocytes. We endow some of the individual synapses with “effective astrocyte ensheathment” by altering their synaptic parameters accordingly, and study the resulting changes in network dynamics.

We start in this work by analyzing simulations of the microscale DiRT (Diffusion with Recharging Traps) model for different levels of astrocyte protrusion into the synaptic cleft. Interestingly, this model finds a strong negative linear relationship between protrusion depth and the strength and time constant of the ensheathed synapse. In other words, astrocyte ensheathment leads to weaker and faster synaptic communication across neurons. We then utilize this linear relationship and implement this result into a series of large-scale spiking models similar to those studied extensively in [2]. This implementation transforms a homogeneous network, a network with uniform synaptic parameters across populations, into a heterogeneous network, where the parameter values for an individual synapse depend explicitly on its level of astrocyte ensheathment. We investigate the impacts this novel heterogeneity has on fundamental states of the network, such as its firing rate, pairwise correlation, and stability of its dynamic states. We demonstrate that, depending on key parameters, such as the number of synapses ensheathed and the strength of this ensheathment, astrocytes can push a non-spatial network from a balanced, asynchronous firing regime into a highly synchronous one. Consistently, we then show that when neuronal assemblies are embedded into our non-spatial network, astrocyte ensheathment can enhance correlations within populations. Finally, broadening our network architectures to include models with spatially structured connections, we find that these results readily extend, namely that the spatial correlation patterns sharpen (show stronger correlation in more spatially-restricted regions) in the presence of astrocytes.

Overall, our results indicate that our understanding of what dynamical modes can be expected from a particular network of neurons needs to be reevaluated in light of synaptic heterogeneity induced by astrocyte ensheathment. Specific ensheathment-induced parameter modification for individual synapses will need to be expanded as we learn more about the effects of the synaptic ensheathment from the detailed models. In the meantime, this framework can easily be applied to other dynamic networks in other contexts using our “effective ensheathment” approach.

## Models

### Microscale model of neurotransmitter diffusion

We use the diffusion with recharging traps (DiRT) model [33, 34] to investigate the number of neurotransmitters that interact with receptors found on the postsynaptic terminal vs. those that diffuse out of the synaptic cleft and interact with the ensheathing astrocyte. We consider a similar idealized synapse as this previous work, taking the cleft to be a two-dimensional domain

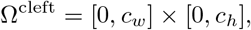

where *c*_*w*_ is the width of the cleft, and *c*_*h*_ is the height. *N*_rec_ postsynaptic receptors of equal size are located along the postsynaptic density

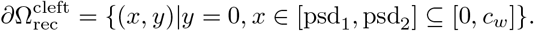

The boundary of the ensheathing astrocyte is taken to be

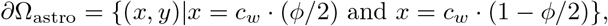

where *ϕ <* 1 denotes the fraction of the synaptic cleft blocked due to astrocyte protrusion (Fig 1A). As a result, stronger astrocyte ensheathment corresponds to larger values of *ϕ*. Note that we also allow for *ϕ* to be negative (i.e., there exists space between the astrocytes and the synaptic cleft), in which case the domain outside of the cleft is referred to as the extracellular space Ω^extra^.

**Fig 1.**
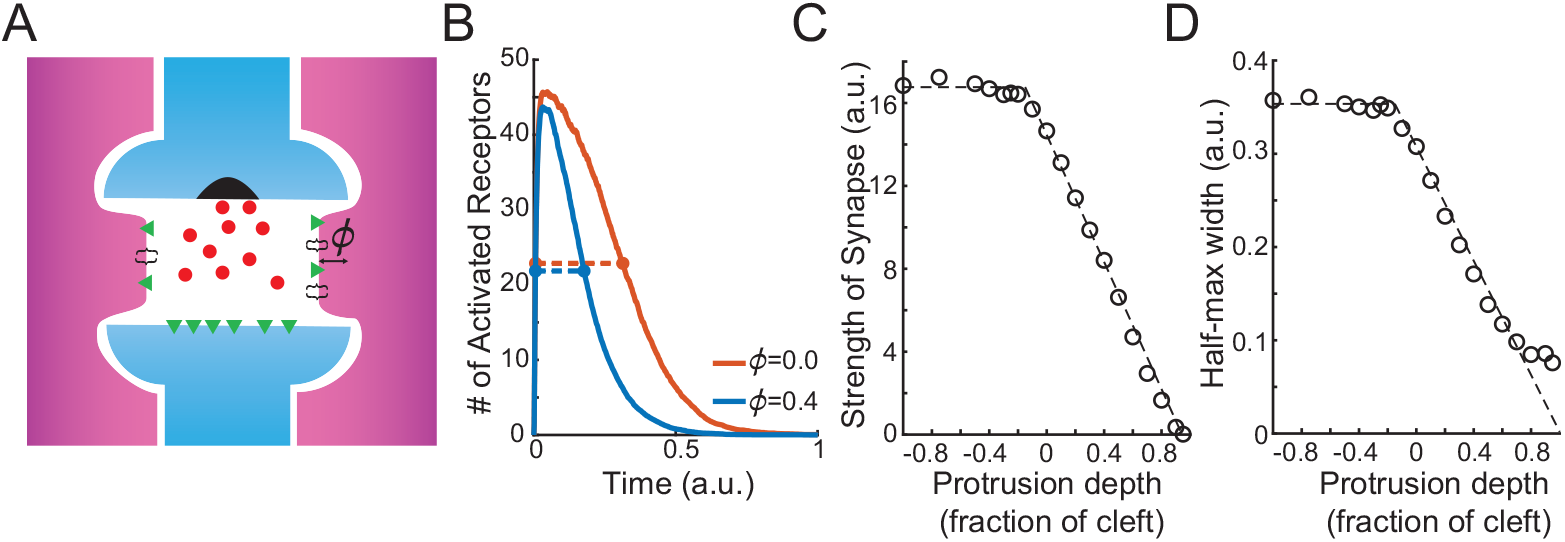
Model of astrocyte ensheathment. A: Schematic of astrocyte (purple) ensheathment of a synapse (blue). The astrocyte is protruding into the synaptic cleft by amount *ϕ*. B: Time course of receptor activation in a diffusion with recharging traps simulation for different depths of astrocyte protrusions (red: *ϕ* = 0, blue: *ϕ* = 0.4) of the neighboring astrocyte, with the half-max widths denoted (dashed lines). C: Strength of synapse, defined as the area under the curve of activation time course, and D: half-max width as a function of protrusion depth. Dashed lines show a piecewise linear fit to the simulated data.

The simulation begins with *N*_NT_ neurotransmitters being released at (*c*_*w*_*/*2, *c*_*h*_).

While in the cleft or in the extracellular space, the neurotransmitters diffuse according to

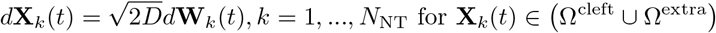

where **X**_*k*_(*t*) denotes the location of the neurotransmitter, **W**_*j*_(*t*) denotes independent Wiener processes, and *D* is the diffusion coefficient. The receptors along the postsynaptic terminal are taken to be partially absorbing, meaning the probability of a particle being absorbed upon making contact is

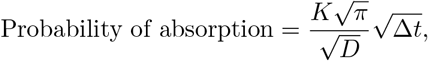

where Δ*t* is the time step of the diffusion model and *K* is the absorption rate. After successfully absorbing a neurotransmitter, the receptor becomes activated and switches to a transitory reflecting state and is unable to bind additional molecules. The time in this transitory state is taken to be exponentially distributed with mean *τ*_*r*_ *>* 0. After this ‘recharge’ time, the receptor switches back to its partially absorbing state. The astrocyte boundary is taken to be perfectly absorbing, with no recharge needed between captures. We define the synaptic time course in this framework as the number of active receptors over time (Fig 1B), and the strength of the synapse to be the area under this curve. All parameter corresponding to this microscale model can be found in Table 1.

**Table 1.** Default parameter value for the DiRT model (arbitrary units).

| Parameter | Default value | Description |
| --- | --- | --- |
| $N_{\text{NT}}$ | 1000 | number of neurotransmitters released |
| $N_{\text{rec}}$ | 50 | number of postsynaptic receptors |
| $\tau_r$ | 0.1 | mean receptor recharge time |
| $K$ | 1 | absorption rate |
| $D$ | 1 | diffusion coefficient |
| $c_w$ | 1 | cleft width |
| $c_h$ | 0.1 | cleft height |
| $[\text{psd}_1, \text{psd}_2]$ | [0.25 0.75] | postsynaptic density location |
| $\phi$ | -1 to 0.95 | fraction of the cleft blocked by protruding astrocyte |
| $\Delta t$ | 0.00001 | diffusion time step |

### Exponential integrate-and-fire neuron

Throughout this paper, we examine the effects that astrocyte ensheathment has on the network dynamics of exponential integrate-and-fire (EIF) neurons (similar to those used in [2]),

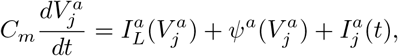

where 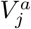 denotes the membrane potential for neuron *j* in population *a* (= *e, i*). The first two terms on the right-hand side correspond to the leak

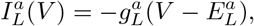

and spike-generating current

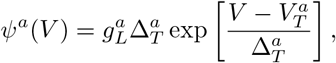

respectively. The third term represents the synaptic input currents, and is given by

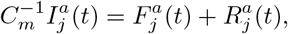

where 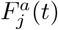 represents the feedforward inputs and 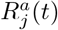 are the recurrent inputs. We consider recurrent inputs of the form

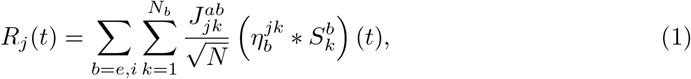

where 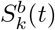 is the spike train, 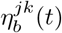 represents the synaptic kinetics and takes the following form

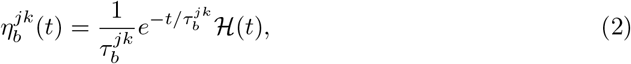

and ℋ (*t*) is the Heaviside function.

### Network connectivity and feedforward input

The 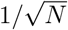 scaling of the synaptic weights 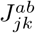 in Eq 1 highlights that we are in the regime of strong recurrent coupling, a key feature of the so-called tightly balanced network [35, 36]. In this regime, if the firing rates of the inhibitory neurons are kept fixed, the strong coupling across excitatory neurons leads to amplification and unstable dynamics [37]. However, if recurrent connections with inhibitory neurons are sufficiently strong and fast, inhibition can effectively track and balance recurrent excitation, leading to asynchronous firing across the network [2, 36–40]. Such asynchronous firing behavior is robust to parameter choices, and is observed across networks with different connectivity rules. The focus in this work is to investigate how the dynamics established by this strong coupling might be modulated by the ensheathment of synapses by astrocytes. To do this, we consider both a randomly-connected and a spatially extended recurrent layer defined by connectivity rules previously considered in this literature. Unless otherwise noted in the figure captions, the parameter used for each network can be found in Table 2.

**Table 2.**
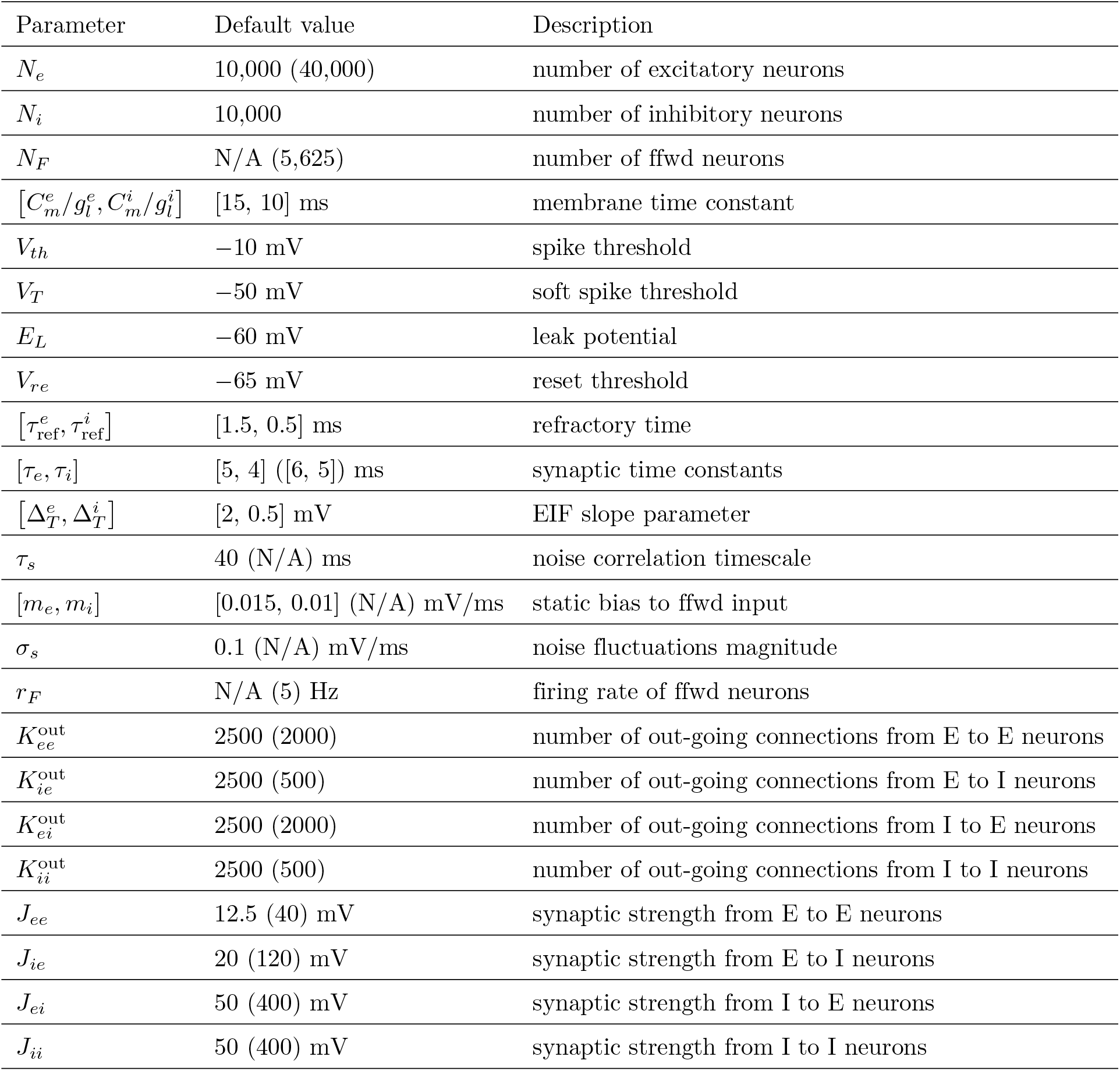
Default parameter value for the non-spatial and spatial model (spatial model parameters in parentheses if different).

### Non-spatial model

In the case of the non-spatial model, for each postsynaptic neuron in population *b*, we randomly and uniformly chose 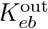 excitatory and 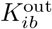 inhibitory presynaptic neurons in the network for it to be connected to (i.e., we fixed the number of outgoing connections). The random network receives feedforward inputs as smoothly varying input biases that can target specific subpopulations. Specifically, the feedforward input to each neuron was given by the sum of an input bias and a smoothly varying signal, 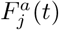

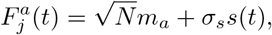

where *s*(*t*) is smooth, unbiased Gaussian noise that is shared across neurons. We take its auto-covariance function to be

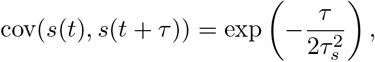

where *τ*_*s*_ sets the correlation timescale and *σ*_*s*_ scales the magnitude of the fluctuations. We consider two cases: 1) all neurons receive the same signal *s*(*t*) (Fig 2A), and 2) half of the neurons receive realization *s*_1_(*t*) and the other half receive *s*_2_(*t*) (Fig 2B).

**Fig 2.**
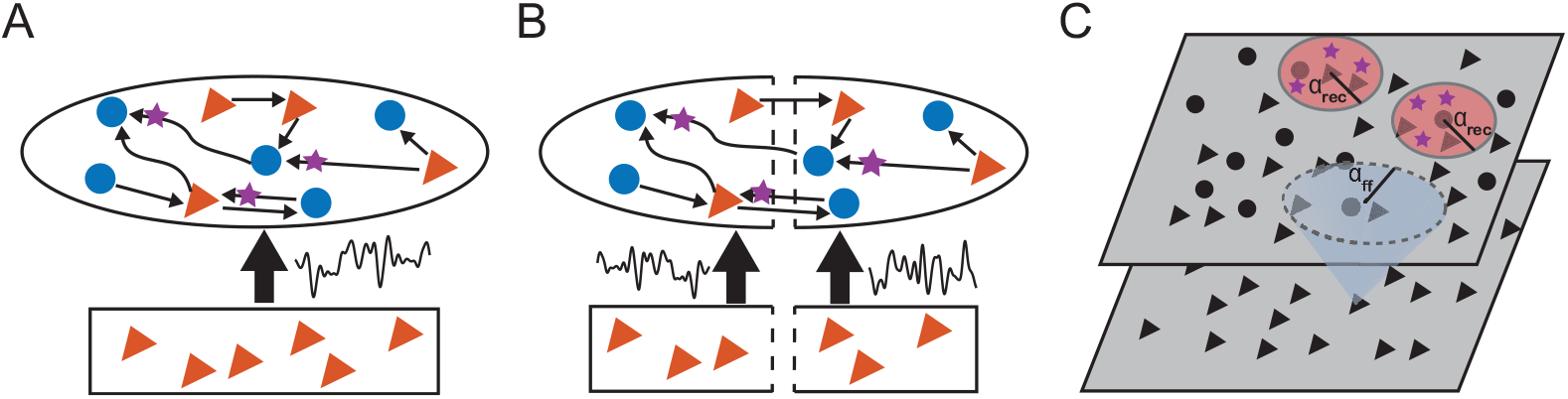
Network schematics. We consider networks of excitatory (red triangles) and inhibitory (blue circles) neurons with different connectivity rules. Individual synapses can be ensheathed by an astrocyte (purple stars). A: One population, non-spatial network where all neurons receive the same Gaussian noise *s*(*t*). B: Two population, non-spatial network where the neurons are randomly split to either receive *s*_1_(*t*) or *s*_2_(*t*). C. Spatial network on a periodic domain, where the recurrent connections can be ensheathed by astrocytes.

To model the spatial network, we arranged *N*_*e*_ excitatory and *N*_*i*_ inhibitory neurons on a uniform grid covering a two-dimensional square domain (Fig 2C). Similarly, the feedforward layer consisted of *N*_*F*_ excitatory neurons uniform grid covering a square that is parallel to the recurrent network. We then normalized each grid such that the domain lied in the unit square Γ = [0, 1] × [0, 1]. Neurons were connected randomly and the probability that two neurons were connected depended on their distance measured periodically on Γ. We used the same algorithm presented in [2] to form the connections. Given a presynaptic neuron at coordinates *y* = (*y*_1_, *y*_2_) in population *b* and a postsynaptic neuron at *x* = (*x*_1_, *x*_2_) in population *a*, this algorithm creates a network where the expected number of synaptic contacts from *b* to *a* is given by

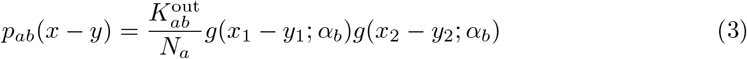

where *g*(*u*; *α*) is a wrapped Gaussian distribution [41]. The *N*_*F*_ feedforward neurons are taken to be Poisson-spiking, so that the feedforward input to the recurrent layer is

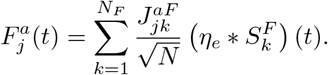

## Results

### Modeling astrocyte ensheathment with an effective astrocyte

We start by investigating how absorption of excess neurotransmitter by astrocytes can shape synaptic time courses. After neurotransmitters are released into the synaptic cleft, they either interact with receptors on the postsynaptic terminal or are absorbed by neighboring astrocytes (e.g., via glutamate transporters). While it is reasonable to assume the depth that an astrocyte protrudes into a cleft impacts the number of neurotransmitters ending up in each destination, it is unclear whether this impact is enough to significantly alter the strength of the transmitted signal. To address this unknown, we make use of the DiRT model [33, 34] to simulate particles interacting with the receptors on this microscale domain (Fig 1A). We find that the protrusion of an astrocyte noticeably decreases the synaptic time course by effectively assisting the cleft in clearing out neurotransmitters (Fig 1B, red: *ϕ* = 0, blue: *ϕ* = 0.4). Further, by defining the strength of the synapse to be the area under the curve of these synaptic time courses, we find that protrusion also causes the synapse to become noticeably weaker. We explore this relationship between the amount of protrusion and synaptic strength in more detail by considering a range of protrusion values (*ϕ* ∈ [−1, 0.95]). In the extreme case where the astrocyte sits significantly away from the cleft (*ϕ <* −0.2), the synaptic strength rests at a constant value (Fig 1C). As the astrocyte approaches and then begins to fill in the cleft (*ϕ* ∈ [−0.2, 0.95]), we see that the protrusion amount and the strength vary linearly, decreasing to zero as the astrocyte completely fills in the extracellular space. This narrative also largely holds true for the half-max width of the synaptic time course (Fig 1D).

Such microscale simulations are computationally intractable to conduct in a large network of spiking neurons, but they provide insight into how one might include “effective” astrocytes in such a model. Namely, a synapse that is ensheathed by an astrocyte should be weaker and faster than an unsheathed synapse. As a result, an ensheathed synapse should have smaller values for *J* and *τ*, which correspond to the synapse’s strength and time constant respectively (see Eqs 1 and 2).

We implement these results of the microscale model in the following way. First, we let *s*_*en*_ ∈ (0, 1) denote the strength of astrocyte ensheathment in the network, where stronger astrocyte ensheathment implies a greater amount of protrusion into the synaptic cleft (i.e., *s*_*en*_ ∼ *ϕ*). Second, for a synaptic connection from neuron *k* to neuron *j*, we let **1**_*jk* ensheathed_ be an indicator function that equals 1 if the *jk* synapse is ensheathed and zero otherwise. We then choose to set this synapse’s strength 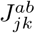 and time constant 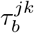 to be

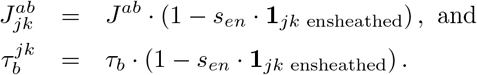

This model captures the key results from the DiRT simulations, namely the synaptic strength 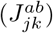 and the half-max width of the synaptic time course (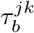 · In(2)) are linearly decreasing functions of the ensheathment strength (Fig 1C,D). Lastly, the probability that an excitatory (inhibitory) synapse is ensheathed is given by 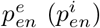. The difference between this and previous network models that accounted for synaptic heterogeneity is that the synaptic strength 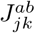 and synaptic time constant 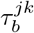 of different synapses are highly variable across the network. Though this adjustment appears deceptively subtle, allowing each synapse to have its own synaptic parameters introduces a level of heterogeneity not seen in previous work.

Utilizing this readily implementable model of astrocyte-neuron interactions, the rest of paper investigates how this heterogeneity introduced by astrocyte ensheathment may modulate the network’s ability to synchronize in several commonly used configurations introduced in Models (Fig 2). Specifically, we consider non-spatial networks receiving one or two smoothly varying input biases and a spatial network driven by a feedforward layer of Poisson-spiking neurons. Since recent experimental evidence suggests that glutamatergic (i.e., excitatory) neurons are preferentially approached by astrocytes [42], we initially focus on exploring the 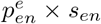 parameter space. Building upon these initial results, we then perform a more in-depth parameter sweep that accounts for the ensheathment of both excitatory and inhibitory synapses.

### Astrocyte ensheathment can break E-I balance, causing synchronous activity

We first consider a non-spatial recurrent layer with each neuron receiving the same smoothly varying signal *s*(*t*) (Fig 2A). Surprisingly, despite each neuron receiving this same input signal, previous work has shown that such a network with homogeneous parameters can exhibit robust asynchronous firing, as long as the network is placed in strongly coupled, or tightly balanced, parameter regime [2, 39]. Placing our EIF network into such a regime reproduces this result, namely a network that exhibits asynchronous spiking (Fig 3A). However, introducing astrocyte ensheathment onto the excitatory synapses broke E-I balance and caused the network to show strong levels of synchrony (Fig 3B and 3C). We emphasize that all three networks in Fig 3 share the same underlying connectivity and receive the same feedforward input. Therefore, the only differences between the networks used in Fig 3B and 3C are the specific synapses that were selected for ensheathment. Together, these results demonstrate that network activity can be driven to synchrony at different times based on the subtle interactions and loss of balance between internal excitatory and inhibitory activity and the external drive.

**Fig 3.**
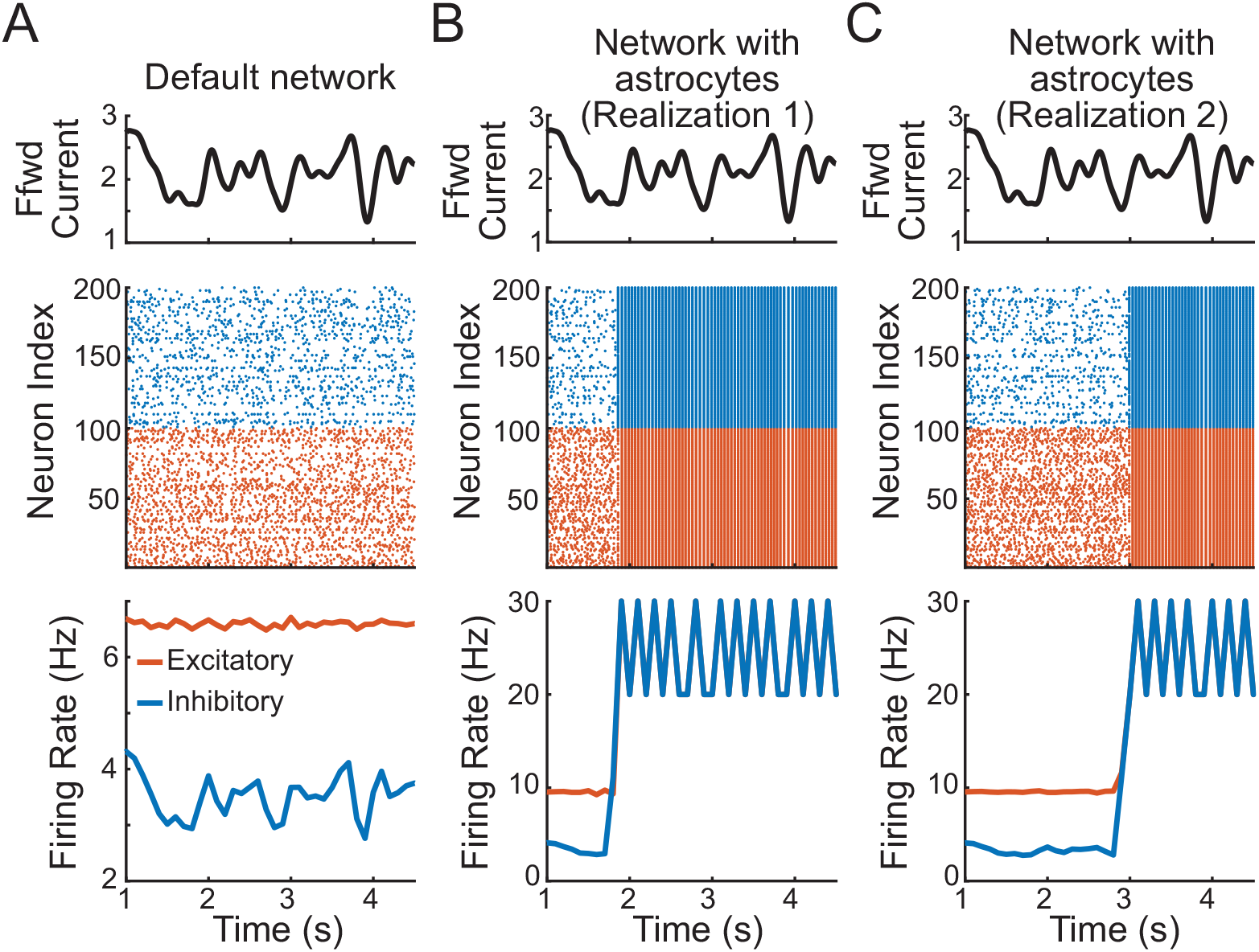
Astrocyte ensheathment can lead to synchronous firing across the network. A: The time course of the input current (top), spiking times of sampled excitatory (red) and inhibitory (blue) neurons from the one population, non-spatial network (middle), and average firing rate or the default network without astrocytes showing asynchronous firing rates (bottom). B and C: same as panel A but for two network with astrocytes realizations, with both networks exhibiting synchronous activity. Ensheathment parameters for B and C: 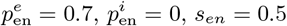.

To investigate what caused the balance state to be lost, we examine the normalized shared current across 500 neurons (Fig 4A; solid: default network, dashed: realization 1 of the network with astrocyte ensheathment from Fig 3B). Initially, we observe that the recurrent activity in both networks tracks and cancels the feedforward input, the crucial characteristic of a balanced network (Fig 4A; gray lines). However, we see that this breaks down at ~1.8 seconds. At this time, the feedforward input into the networks significantly decreases (Fig 4A; black line), and the recurrent activity in the astrocyte network is unable to successfully cancel it. As a result, the balanced state is lost and the network transitions into synchronous firing.

**Fig 4.**
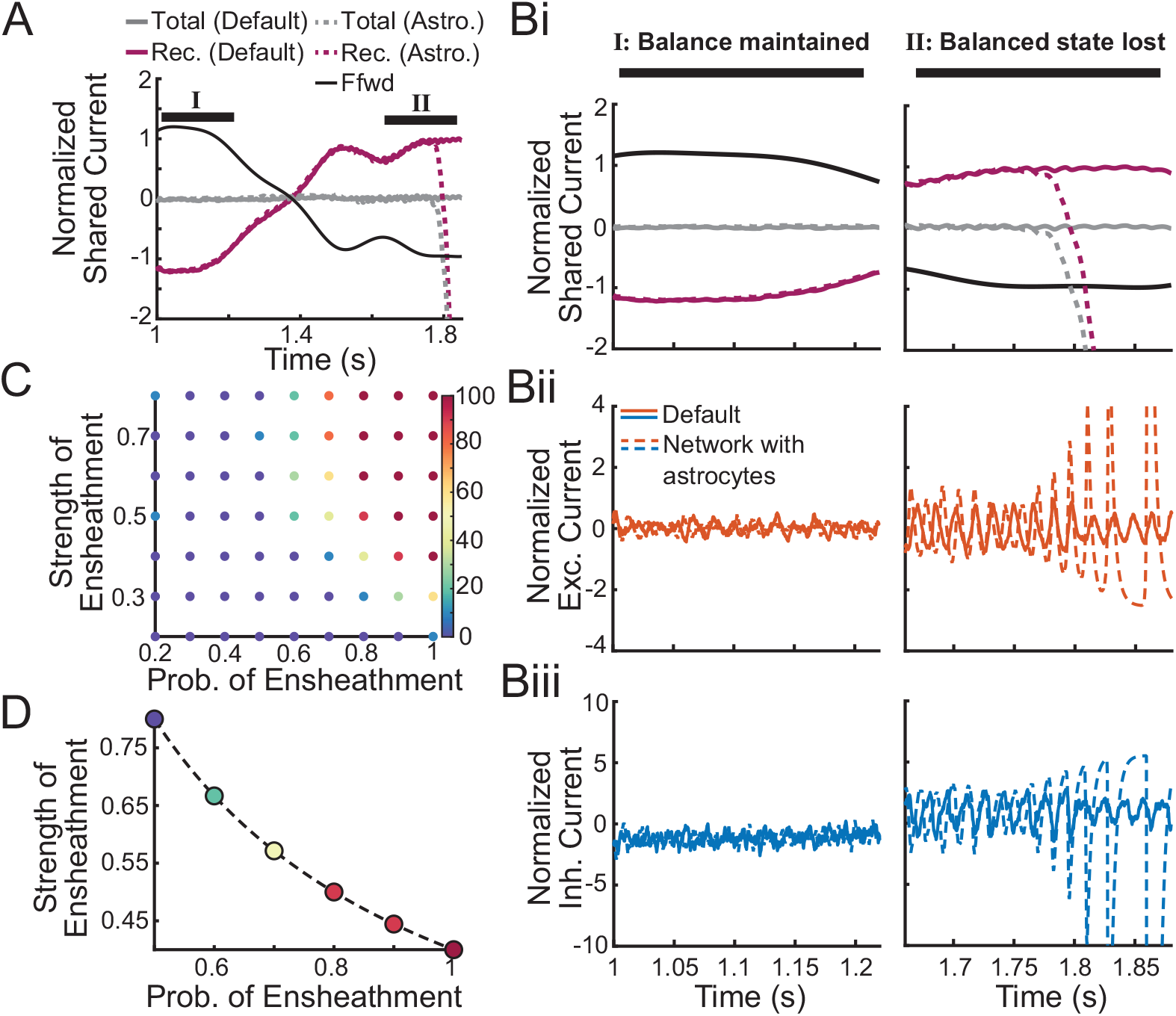
Astrocyte ensheathment can break E-I balance. A: Normalized shared fluctuations in the feedforward (black) and recurrent (purple) synaptic inputs for two one population, non-spatial networks (solid: default; dashed: realization 1 from Fig 3B). Initially, both networks exhibit recurrent dynamics that effectively cancel the feedforward inputs, leading to weak shared fluctuations (gray), until the balance state is lost in the network with astrocyte ensheathment. The curves were computed by averaging the synaptic input currents to 500 neurons, convolving with a Gaussian-shaped kernel (*σ* = 15 ms), subtracting the mean and dividing by the neurons’ rheobase. B: Normalized shared fluctuations in the currents zoomed in on a time window when the asynchronous state is maintained in both networks (left column) and for the time window when the asynchronous state is lost in the network with astrocyte ensheathment (right column). The following currents are displayed: i) feedforward, all recurrent, total, ii) recurrent excitatory, and iii) recurrent inhibitory currents in the default network (solid) and the network with astrocyte ensheathment (dashed). C: Percent of the ten EIF networks that led to synchronous dynamics in 5 seconds of simulation as a function of ensheathment parameters. D: The 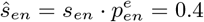 isoline showing that a naïve mean field approximation would fail at capturing this phenomenon. Colorbar is the same as panel C. 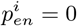 for all panels.

To better understand the mechanisms supporting the asynchronously firing state and, more importantly, what causes the loss of this state in the network with astrocyte ensheathment, we take a closer look at two time windows during this simulation (Fig 4B). During the first time window (I), the balanced state is maintained in both networks. We note that the normalized feedforward current is large and positive, and by necessity, the net shared/recurrent current is negative to compensate (Fig 4Bi). This cancellation of the feedforward input is achieved via small fluctuations in the excitatory currents that are then effectively tracked and balanced by the inhibitory currents accordingly (Fig 4Bii and 4Biii).

However, during the second time window (II), the normalized feedforward current drops, which must be met with an increase in the net shared current. This increase runs the risk of leading to uncontrolled runaway excitation, and as a result, synchronous firing. We see that the default network’s inhibitory currents are able to continue tracking and balancing the excitatory currents, preventing this from occurring.

Meanwhile, the network with astrocyte ensheathment struggles to make these fine tuned adjustments, which leads to growing oscillations in both the excitatory and inhibitory currents. Interestingly, we note that the second realization of the astrocyte ensheathment network (Fig 3C) does not fall into synchrony at this point in the simulation, and instead is able to maintain the balanced state during this drop in the feedforward current (S1 Fig D, center column) due to random differences between realizations (i.e., the specific synapses chosen for astrocyte ensheathment). Nonetheless, it fails later on in the simulation when the feedforward input reaches a more extreme local minimum (S1 Fig D, right column).

To explain this loss of the asynchronous state, we recall that our model of astrocyte ensheathment leads to faster and weaker synapses. As noted previously, the balanced state relies on inhibition to be sufficiently fast relative to excitation [39, 40]. Here, astrocytes are specifically targeting excitatory synapses, so the net effect is known: a subset of excitatory synapses throughout the network now have faster synaptic dynamics. As these simulations then suggest, their inhibitory counterparts are unable to effectively track their activity and the network experiences runaway excitation as the balanced state is lost. While consistent with previous work, our results show that adjusting the synaptic kinetics in only a *subset* of neurons is sufficient for the entire network to enter into a state of synchronous firing.

We now explore how the parameters underlying astrocyte ensheathment (i.e., the probability 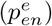 and strength (*S*_*en*_) of ensheathment of excitatory synapses) determine the size of subset needed to lead to synchronization and whether a mean-field approximation can be used to average across this heterogeneity and capture this effect. We consider ten default EIF networks, each with a different input signal *s*(*t*) and no astrocyte ensheathment. These ten networks all exhibited asynchronous spiking solutions for the full length of the trial (taken to be 5 seconds). We then ask whether or not these same networks synchronized in the presence of astrocyte ensheathment. Discretizing the parameter space 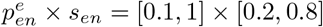 into seventy parameter sets, we observe that all ten networks eventually exhibit synchronous behavior as 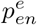 and *s*_*en*_ increase (Fig 4C; red dots in upper right corner). However, we also observe that there is not a sharp transition point from blue (all networks remain in the asynchronous state) to red (all networks become synchronous). This suggests that for intermediate values of 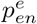 and *s*_*en*_, random differences in the connectivity diagram and the locations of astrocyte ensheathment are enough to prime some networks to tend toward synchrony over others.

These results highlight the shortcomings that a naïve mean field theory would have in capturing the dynamics brought on by this type of synaptic heterogeneity. More specifically, one might be tempted to simply average out the effect of astrocyte ensheathment across the network and adjust the strength of all synapses to be

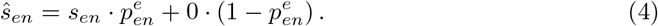

However, by considering the outcomes across the 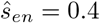 iso-line (Fig 4D), the heterogeneous networks may rarely exhibit synchrony (low 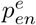, high *s*_*en*_) or frequently (high 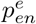, low *s*_*en*_).

### Astrocyte ensheathment can enhance correlations already present within a network

The previous section explored how astrocyte ensheathment of excitatory synapses affects the dynamics of a balanced network operating in the asynchronous regime. Now we examine a network with the same non-spatial connectivity, but the neurons are split into two subpopulations, where each subpopulation receives its own feedforward input (Fig 2B). As previously shown in [2], this change in feedforward inputs leads to a dramatic shift away from the asynchronous state, with positive correlations arising among neurons within the same population and negative correlations across populations. As a result, the correlation distribution becomes strongly binomial (Fig 5A,B; solid lines). While we expect the results of the previous section to carry through, namely the loss of stability for the strong ensheathment of excitatory synapses, it is unclear how the heterogeneity introduced by more moderate levels of ensheathment will modulated these new correlations inherently found in the network.

**Fig 5.**
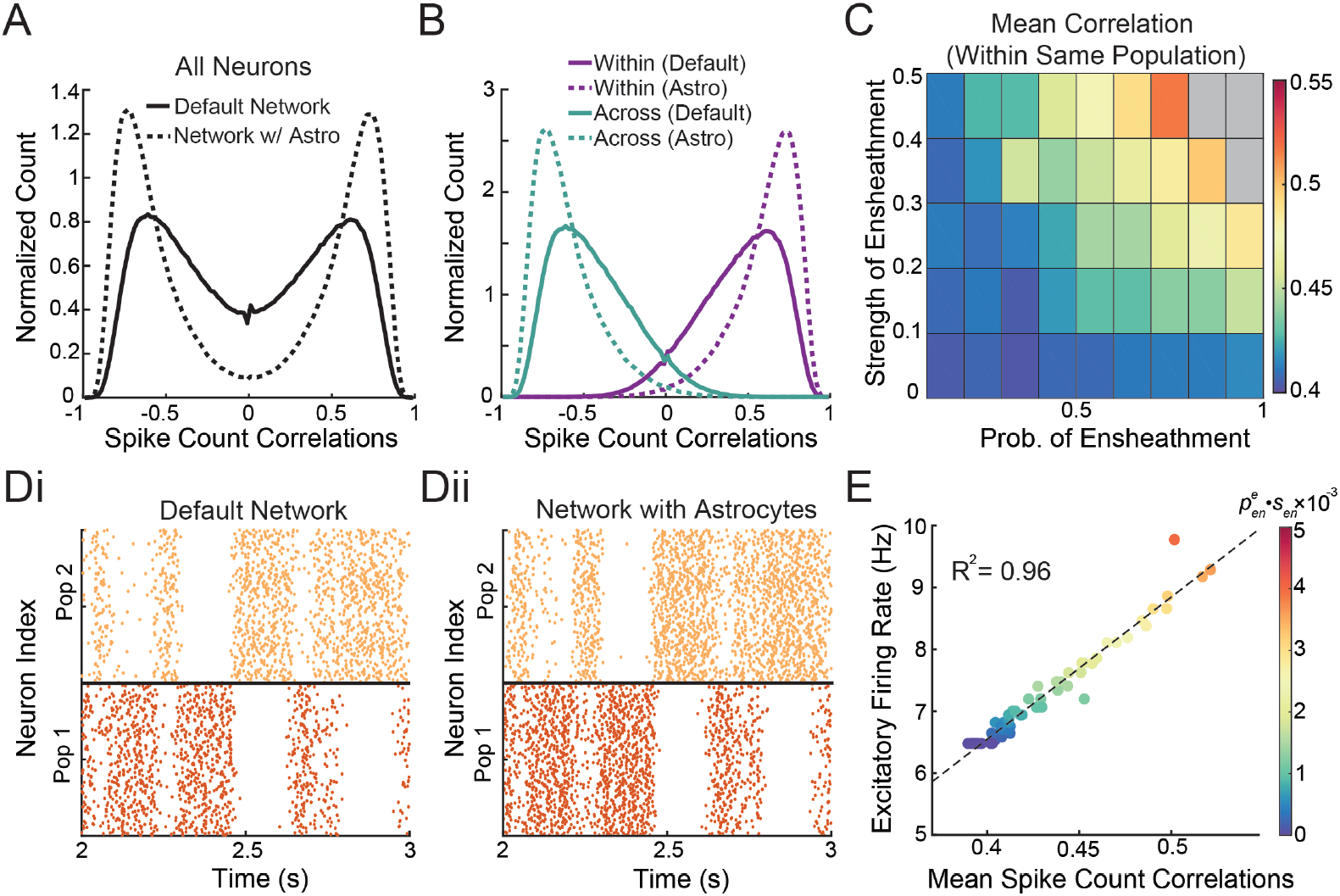
Astrocyte ensheathment can lead to greater correlations. A: Normalized histogram of pairwise spike count correlations between 1000 neurons for the default two population, non-spatial network (solid) and the same network with astrocytes (dashed), with parameters 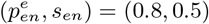. B: Same as panel A, except the pairwise spike count correlations calculated within and across each populations. C: Mean pairwise spike count correlations within the same population as a function of ensheathment parameters. Gray regions denote areas where the network lost stability. D: Raster plots for the two populations for the i) default network and ii) network with astrocytes 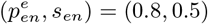. E: Mean spike count correlations plotted against the excitatory firing rate for different astrocyte conditions (the color of dots corresponds to 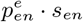 for that simulation). 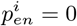 for all panels.

Setting 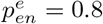 and 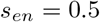, we find that the binomial distribution of spike count correlations present in the default network becomes more extreme, with an increase in the positive correlation among the same population and a decrease in the negative correlations across populations (Fig 5A,B; dashed lines). This trend is true in general: as 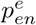 and *s*_*en*_ increase, the correlations among a single population also increase (Fig 5C), while the correlations across different populations decrease (figure not shown). As anticipated, we also find that the network stability is loss for more extreme values of astrocyte ensheathment (Fig 5C, gray boxes).

To gain additional insight into how astrocytes are modulating the spiking behavior of the network, we compare the raster plot of the default network to a network with astrocytes, where both networks receive the same feedforward inputs (Fig 5D). As expected from the presence of negative correlations across populations (Fig 5B), the two populations show clear competitive dynamics. Interestingly, the network with astrocyte ensheathment shows a much more subtle modulation of network dynamics than our previous results. In fact, it is clear from these raster plots that key quantitative traits, such as the times the dominate spiking populations switches, are largely conserved across the two conditions.

One observable difference between the two networks is the population firing rate: both populations in the network with astrocyte ensheathment appears to have higher firing rates. Previous work [43] suggests that this difference in firing rates could be sufficient to explain the increase in correlations. Indeed, we find a strong positive relationship (R^2^ = 0.96) between the spike count correlations and the excitatory firing rate, with higher firing rates arising with stronger astrocyte ensheathment (Fig 5E; color corresponding to 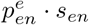). This strong relationship between firing rate, correlations and astrocyte ensheathment suggest that unlike the last network configuration, a mean-field model utilizing the parameter defined in Eq 4 could be used to capture the effect of astrocytes on the neuronal dynamics. See Discussion for additional considerations regarding a mean-field approach.

### Spatial correlations are enhanced by astrocyte ensheathment

In the previous two sections, we showed that the ensheathment of excitatory synapses by astrocytes strongly modulated the underlying correlations across neurons in a non-spatial network. However, it is unclear the impact this ensheathment may have on networks with more realistic connectivity. To pursue this, we now implement our “effective” astrocytes in a spatial model (Fig 2C). Specifically, we arrange the recurrently connected neurons in a two dimensional grid, with a Gaussian connectivity profile (see Eq 3), the spread of which is given by parameter *α*_rec_. Unlike the previous sections, the feedforward inputs are provided by neurons that are Poisson-spiking. These neurons also lie on a two-dimensional grid and make connections based on a Gaussian connectivity profile with spread parameter *α*_ff_. We continue to consider astrocyte ensheathment of only recurrent excitatory synapses.

In agreement with previous work [2], we find that when the spatial profile of recurrent connections is less than the feedforward (i.e., *α*_rec_ *< α*_ff_), the default network exists in a spatially asynchronous state, with small pairwise spike count correlations between neurons of any distance (Fig 6A, left and 6B, black). In fact, [2] proves that the asynchronous state requires *α*_rec_ *< α*_ff_. However, by introducing astrocyte ensheathment to this model (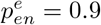 and *s*_*en*_ = 0.5), we find that the asynchronous state can be broken without changing the spatial profiles of connectivity. Specifically, we observe strong positive correlations arising between nearby neurons and negative correlations from those farther away (Fig 6B, orange), resulting in tight clusters of spiking neurons (Fig 6A, right). We note these tight clusters do not persist across time (S1 Video).

**Fig 6.**
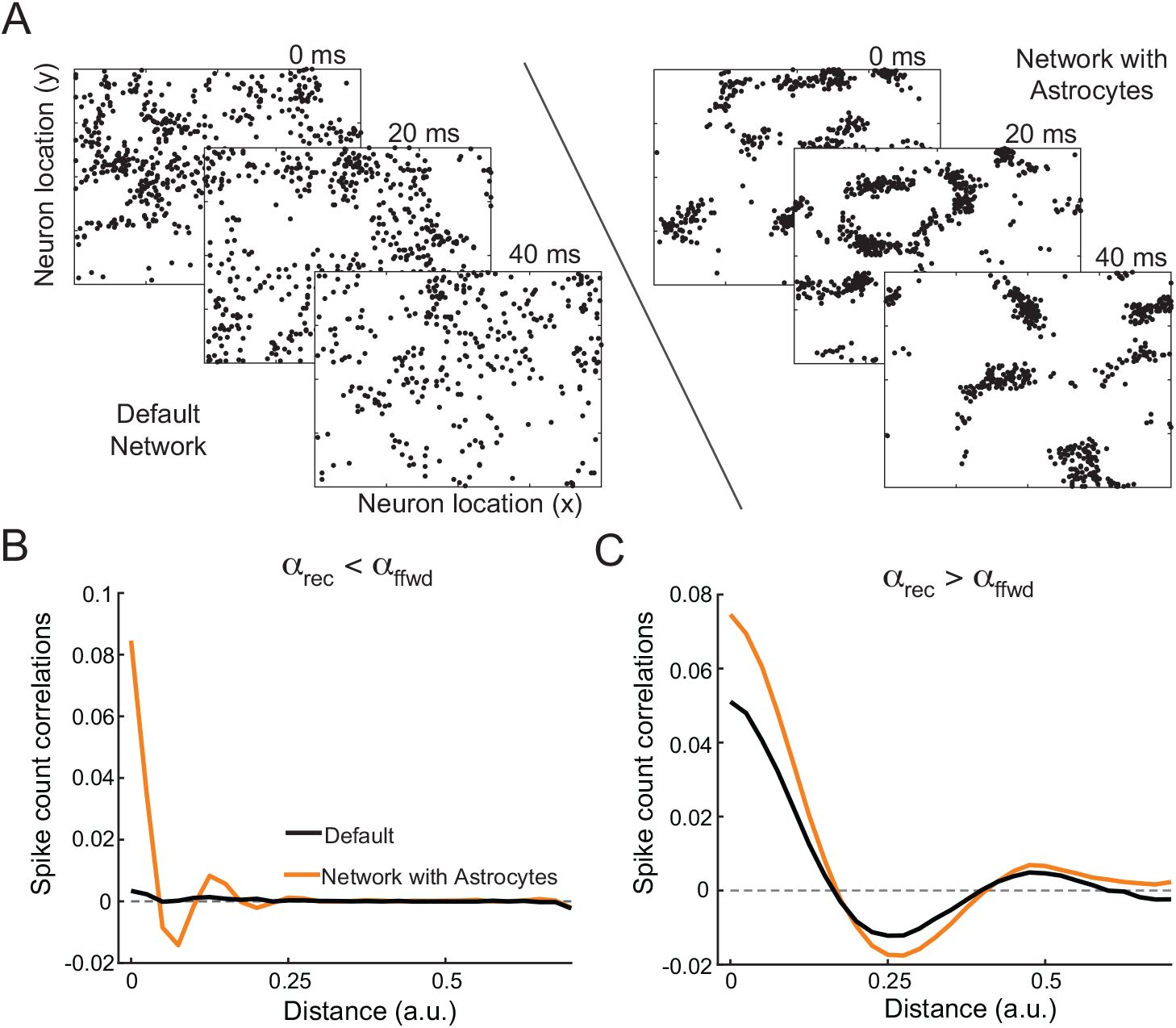
Astrocyte ensheathment can lead to greater spatial correlations. A: Spatial raster plots at three 20 msec time points for the default network (left) and network with astrocyte ensheathment (right). Parameters were chosen such that the default network exhibited no spatial correlations (*α*_rec_ = 0.05 and *α*_ffwd_ = 0.1). Ensheathment parameters are 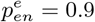 and *s*_*en*_ = 0.5. B: Spike count correlation plots as a function of distance between neuron pairs corresponding to the parameters in panel A for the default (black) and network with astrocytes (orange). C: Same as panel B, except with a default network placed in the spatial correlation regime (*α*_rec_ = 0.2 and *α*_ffwd_ = 0.1). Here, the ensheathment parameters are 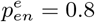 and *s*_*en*_ = 0.5). 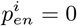 for all panels.

These results mirror our earlier results corresponding to the non-spatial network with a single population of neurons (Fig 3), and arise from similar mechanisms. Specifically, astrocyte ensheathment speeds up excitatory synapses, and as a result, the inhibitory synapses are unable to track and cancel recurrent excitation, leading to a loss of the asynchronous balance state. However, unlike the result for the non-spatial network, the network activity organizes in tight spatial patterns, as opposed the synchronous firing of the entire network.

In further agreement with [2], when the recurrent connection’s spatial profile is larger than the feedforward projections (i.e., *α*_rec_ *> α*_ff_), we find that the default network now exhibits spatial correlations (Fig 6C, black). In this case, astrocyte ensheathment of excitatory synapses does not qualitatively change this behavior, but instead simply enhances these spatial correlations at almost all distances (Fig 6C, orange), with the exception being the distances where spatial correlations are zero. This is true for a range of astrocyte ensheathment parameters: stronger astrocyte ensheathment, either through 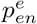 or *s*_*en*_, leads to stronger spatial correlations, while not affecting the spatial footprint of these correlations (S2 Fig). In this case, with the asynchronous state already broken due to the widths of the spatial projections, astrocyte ensheathment of excitatory synapses simply amplifies the inability of inhibitory neurons to effectively track and cancel recurrent excitation across all spatial scales.

### Astrocyte ensheathment of both excitatory and inhibitory synapses can boost stability and within-population correlations

Thus far, we have explored the effects on network dynamics when excitatory synapses are targeted by astrocyte ensheathment. We found that across networks with different connectivity rules and feedforward input, the neuronal dynamics shifted towards a more correlated regime. However, despite evidence suggesting that excitatory synapses are preferentially targeted [42], it is important to note that inhibitory neurons can and are ensheathed, and that the level of ensheathment may vary by the particular brain region. For this reason, we now consider astrocyte ensheathment that can target both types of synapses, expanding our parameter search over the larger 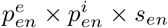 space to examine whether our previous results carry through.

For this, we return to the non-spatial model with two subpopulations that each receive its own feedforward input (Fig 2B). Recall that in this framework, the within population correlations were positive and astrocyte ensheathment of excitatory synapses increased these correlations (Fig. 7A, top left). Across different values of 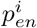, we see that this relationship is not only intact, but amplified: As 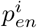 increases, the within population correlations also increase. Further, we note the network’s stability is more robust, with the higher levels of 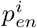 (i.e., ≥ 0.4) lacking the unstable gray regions.

**Fig 7.**
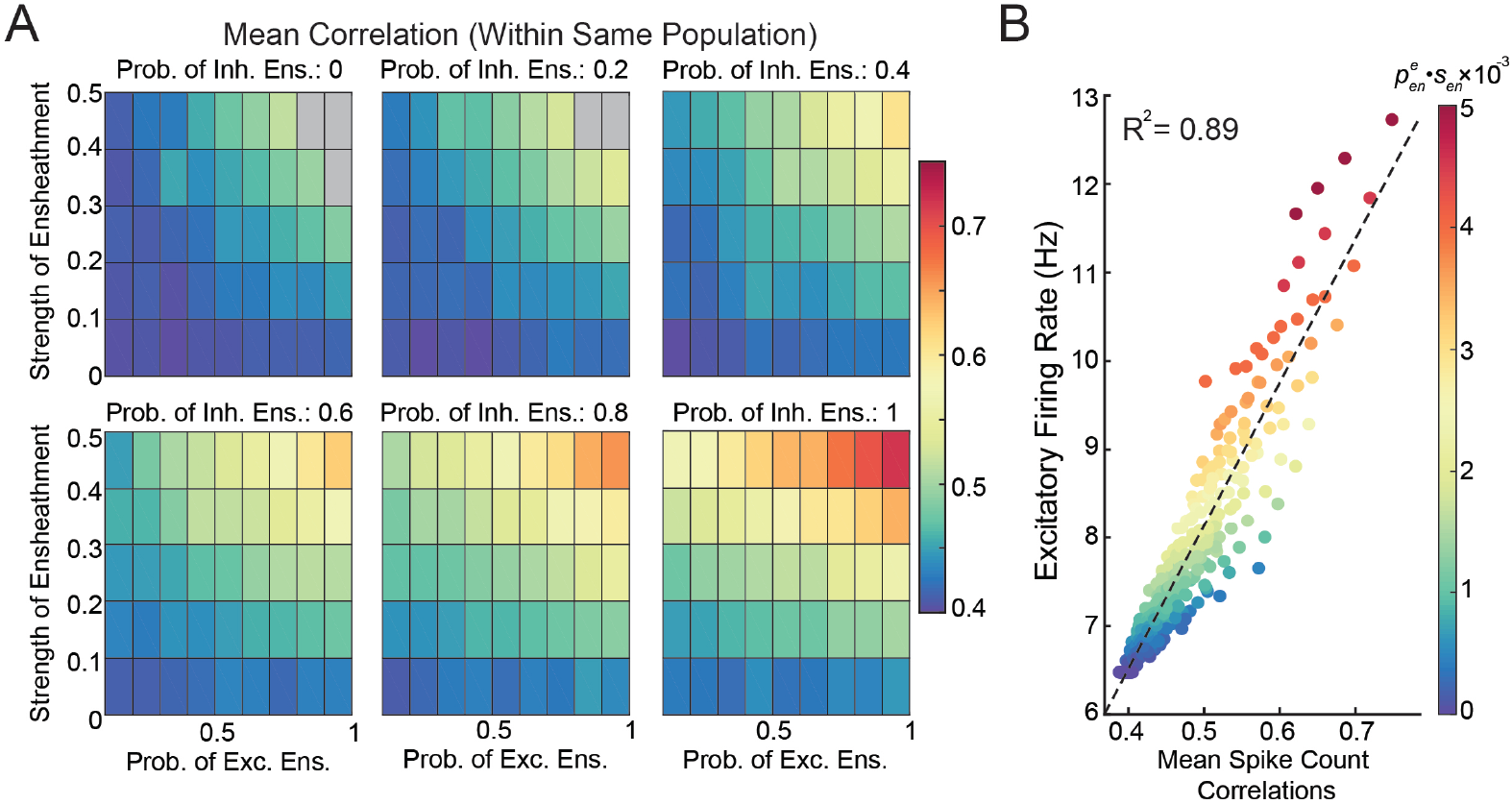
Investigation of neuronal dynamics when astrocyte ensheathment targets both excitatory and inhibitory synapses. A: Mean pairwise spike count correlations within the same population as a function of ensheathment parameters. Gray regions denote areas where the network lost stability. B: Mean spike count correlations plotted against the excitatory firing rate for different astrocyte conditions (the color of dots corresponds to 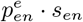 for that simulation).

Lastly, we investigated whether the relationship across correlations, excitatory firing rate, and average ensheathment level of excitatory synapses 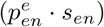 remains. Indeed, we again find a strong linear relationship between correlations and firing rate, and both metrics with a clear correspondence to the strength of astrocyte ensheathment (Fig 7B). In total, these results suggest that while the ensheathment of inhibitory neurons in addition to excitatory do not drastically alter our earlier results, stability and correlations are both reliably enhanced. As a result, this could be a key mechanism utilized by the brain to support high levels of correlation while preventing pathological states from arising.

## Discussion

Through microscale modeling of neurotransmitter diffusion, this work derives an efficient way to implement astrocytes ensheathment of synapses in large scale neuronal networks. Specifically, simulations on the microscale domain of the synaptic cleft suggested that ensheathed synapses are weaker and faster than their non-ensheathed counterparts. This investigation identified the synaptic strength and synaptic time constant as the key parameters to vary to denote the presence of an “effective” astrocyte.

While we have chosen to consider a network of EIF neurons, such parameters are commonly found across different types of recurrently connected spiking networks (e.g., networks of current- and conductance-base neurons, with and without adaptation [37, 40, 44–46]), making the implementation presented here applicable to many network architectures and parameter regimes. We showed this specifically by examining the consequences of including these effective astrocytes in both non-spatial and spatial networks placed in the regime of strong recurrent coupling, which is also referred to as the “tight balanced” regime [36]. In this regime, we uncovered that astrocyte ensheathment can dramatically change firing dynamics, break the balance state, and cause synchronous behavior. While this parameter regime yields a type of inhibition-stabilized network (ISN) [47], meaning the fixed point is stabilized by feedback inhibition, it is unclear whether our results extend to other types of ISN balanced networks (e.g., the “loose balance” regime, also referred to as a stabilized supralinear network [36, 48]) and non-ISN parameter regimes. For example, we were able to explain the synchronous firing in our non-spatial network (Fig 3) as due to the astrocytes targeting and speeding up excitatory synapses, which the inhibitory currents were unable to track and cancel. In these other parameter regimes, this tight canceling does not occur, so it is unclear if similar synchronous firing would arise. Future work will push these lines of inquiry further, made easier in part by the straightforward implementation of our astrocyte framework.

The investigation of the balance state as a function of synaptic time courses was also completed recently in [40], but our work differs in a few notable areas. First, the adjustment of the synaptic time courses here arose from the implementation of astrocyte ensheathment around individual synapses. This resulted in a heterogeneous network, which behaves qualitatively differently than its homogeneous network counterpart found by a naïve mean field (see Fig 4). Second, we observed spatial patterns emerge in the *α*_rec_ *< α*_ff_ regime when astrocyte ensheathment was present, showing that this condition, as derived in [2], is a necessary condition for the balance state, but not a sufficient one. Meanwhile [40] only considered homogeneous networks in the *α*_rec_ *> α*_ff_ regime.

We also note that the heterogeneity introduced by astrocyte ensheathment considered here is significantly different than other models that consider heterogeneous parameters. For example, previous works have implemented heterogeneity in connection strengths by having the 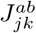’s be randomly distributed according to the normal distribution 𝒩 (*J, σ*) [18–20], resulting in only small variations of parameters across the network. Here, the synaptic parameters corresponding to ensheathed and unsheathed synapses are pulled from entirely distinct distributions, specifically

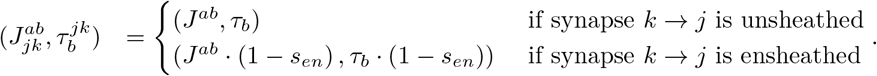

In the future work, it would be important to extend this heterogeneity even further, either by considering distributions for the synaptic parameters as done in previous works, or by considering a distribution on the ensheathment parameter *s*_*en*_. This would allow various degrees of astrocyte ensheathment across a single network and potentially across different populations (i.e., define 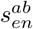 as the strength of ensheathment from population *b* to *a*). In addition, since astrocytes segregate the space into non-overlapping control domains, which can also be disrupted in epilepsy [49]), it would also be interesting and important to perform a more thorough investigation that includes spatial inhomogeneities across all ensheathment parameters for the spatial network. Further, it may also be interesting to consider a time varying component to these ensheathment parameters. In fact, along with increasing the number of postsynaptic receptors [50], long-term potentiation induction has recently been shown to decrease the ensheathment level of astrocytes [51]. While these two mechanisms operate on a much slower timescale as considered in this work, as shown here and in our previous work [33], they would work together to strengthen and slow synapses. Including all of these additional forms of heterogeneities, spatially and temporally, would lead to a more robust understanding of how astrocytes are modulating network dynamics via ensheathment.

Additionally, we found that it is non-trivial to capture this novel form of network heterogeneity in a mean-field approximation, complicating the extension of this framework to be included in rate-based models of average population dynamics. Specifically, when we considered astrocyte ensheathment of excitatory synapses, we showed that a naïve mean-field theory (see Eq 4) would fail to capture key qualitative dynamics of the full model (Fig 4D). On the other hand, when we allowed astrocytes to equally ensheathed both excitatory and inhibitory synapses, we found a tight relationship between correlations, firing rates, and astrocyte ensheathment (Fig 5E), suggesting that a mean-field approximation in this parameter regime could be possible. So while a mean-field approach could work in the latter case, it would be interesting future work to investigate how robust such an approximation would be in the presence of symmetry breaking changes in parameters, either in the probability and/or strength of ensheathment.

Lastly, our simplified model of astrocyte-neuron interactions at the synaptic level only accounts for the ability of astrocytes to assist in the clearance of neurotransmitters from the synaptic cleft by acting as an effective buffer and transporter [13, 52], but other astrocyte-neuron communication pathways are likely to play a role in modulating neuronal dynamics. For example, neurotransmitters may also activate receptors found on astrocyte membranes (e.g., mGluR5), leading to the activation of calcium pathways [53, 54]. While the downstream effects of this pathway are still being investigated, evidence suggests that it may impact other aspects of neuronal signaling not considered in this work, such as the overall excitability of the network by either adjusting the resting potential of neurons [55] or by modifying neurotransmitter profile in the extracellular space through the direct release of gliotransmitters [15] (but see [56–58]). Here, we show the unanticipated result that this calcium pathway is not needed for astrocytes to strongly tune network dynamics.

Astrocytes are also known to have a role in restoring the resting membrane potential of nearby neurons and preventing hyperexcitability through the removal of potassium via Kir4.1 channels [59]. It is therefore possible that increased ensheathment, as considered in this work, might result in improved potassium removal by bringing astrocyte Kir4.1 closer, which may counteract to some extent the tendency to synchronize. On the other hand, rapid reduction of extracellular potassium would also render glutamate removal more efficient [60, 61] making the synchronization that we observe here even more likely. Lastly, our model suggests that tight ensheathment by astrocytes would drastically decrease the amount of neurotransmitters that ‘spillover’ into the extrasynaptic space, which is in alignment with experimental observations [51, 62]. Since such spillover is known to allow for cross-talk between neurons, it is another mechanism that could set the excitability of the network [63, 64]. Future work should seek to connect these additional communication pathways to the ensheathment interaction investigated here to further advance our understanding of how astrocyte-neuron interactions modulate spiking dynamics across brain regions.

## Supporting information

**S1 Video. Video of spatial raster plots showing tight clusters of spiking neurons in the network with astrocytes and asynchronous spiking in the default network**. Will be made available upon acceptance for publication.

**S1 Fig.**
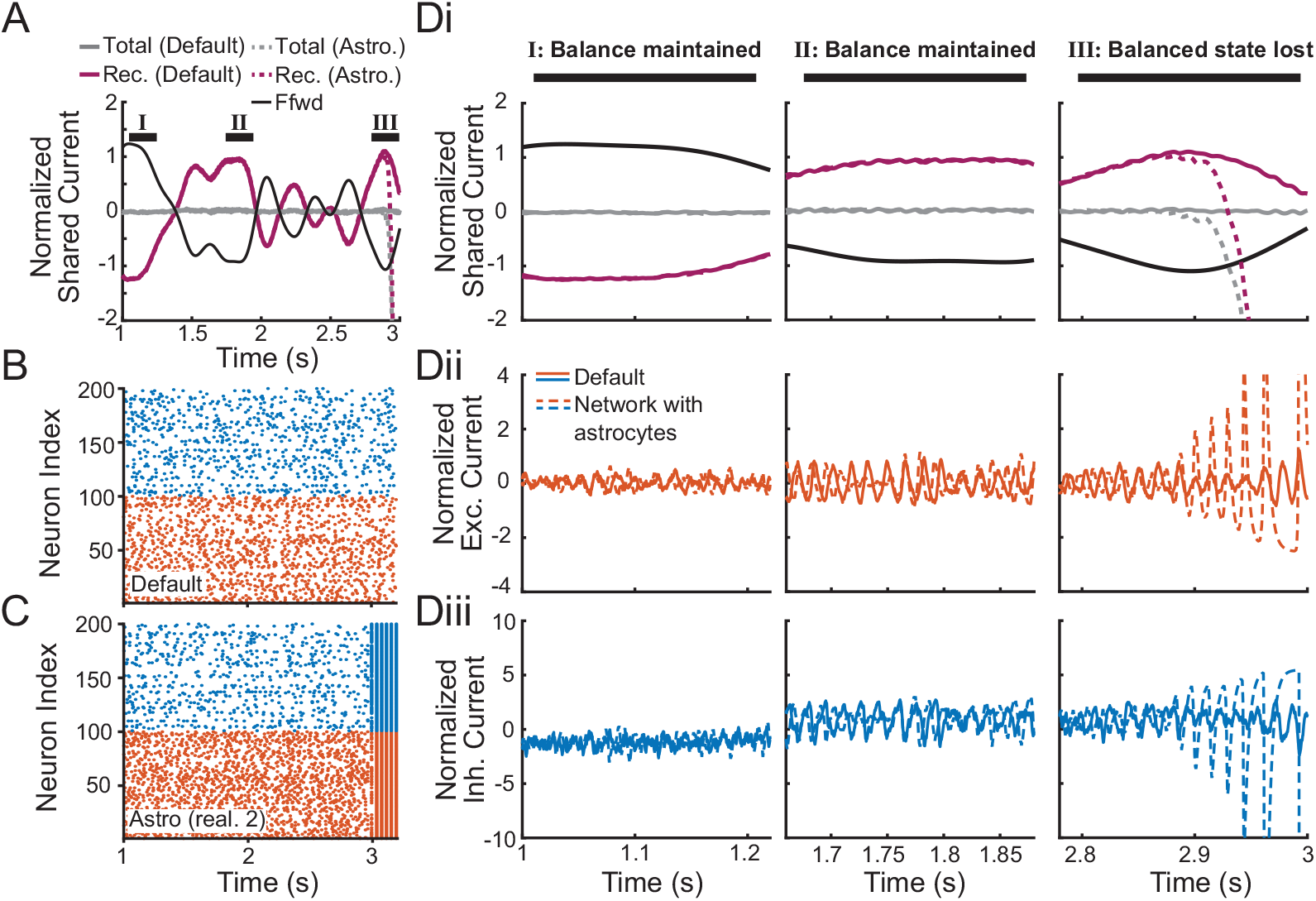
Broken E-I balance in Realization 2. A: Normalized shared fluctuations in the feedforward (black) and recurrent (purple) synaptic inputs for two one population, non-spatial networks (solid: default; dashed: realization 2 from Fig 3C). B,C: Spiking times of sampled excitatory (red) and inhibitory (blue) neurons from the default network and network with astrocyte ensheathment, respectively. D: Normalized shared fluctuations in the currents zoomed in on time windows when the asynchronous state is maintained in both networks (left and middle columns) and for the time window when the asynchronous state is lost in the network with astrocyte ensheathment (right column). The following currents are displayed: i) feedforward, all recurrent, total, ii) recurrent excitatory, and iii) recurrent inhibitory currents in the default network (solid) and the network with astrocyte ensheathment (dashed). Same parameters as Fig 3-4.

**S2 Fig.**
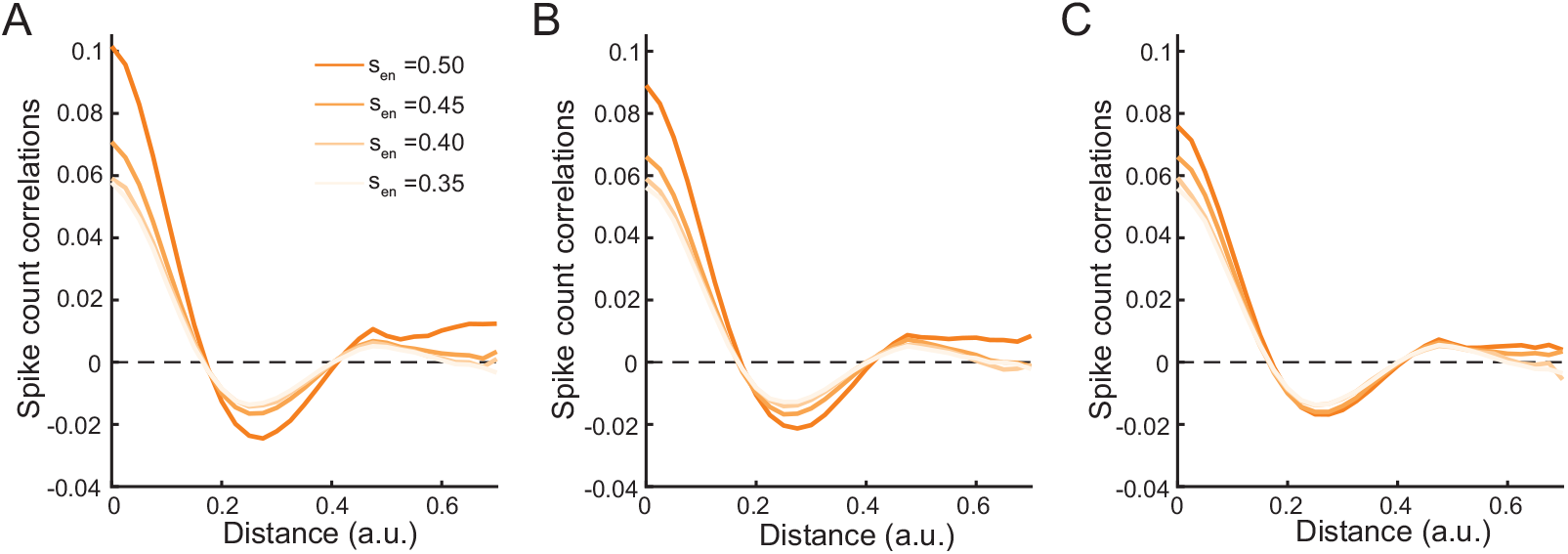
Spatial correlations as a function of ensheathment parameters. Spike count correlations as a function of distance for different probabilities of ensheathment (A: 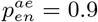, B: 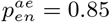, C: 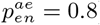) and ensheathment strengths (*s*_*en*_ ∈ [0.35, 0.5]), where the default network exhibits spatial correlations (*α*_rec_ = 0.2 and *α*_ffwd_ = 0.1).

## Data availability

The code used to generate the main figures in the manuscript is available on the GitHub database (https://github.com/gregoryhandy/Neuronal_Net_with_Astro_Ensheathment).

## Acknowledgments

G.H. would like to acknowledge support from the Swartz Foundation Fellowship for Theory in Neuroscience and the Burroughs Wellcome Fund’s Career Award at the Scientific Interface. A.B. would like to acknowledge support from National Science Foundation award NSF-DMS-1853673 and the support and resources from the Center for High Performance Computing at the University of Utah.

